# Unraveling the role of γδ T cells in the pathogenesis of an oncogenic avian herpesvirus

**DOI:** 10.1101/2023.12.05.570256

**Authors:** Mohammad A. Sabsabi, Ahmed Kheimar, Yu You, Dominik von La Roche, Sonja Härtle, Thomas W. Göbel, Theresa von Heyl, Benjamin Schusser, Benedikt B. Kaufer

## Abstract

Marek’s disease virus (MDV) is an oncogenic alphaherpesvirus that causes deadly T cell lymphomas in chickens. MDV is highly cell associated which allows the virus to evade antibody-mediated virus neutralization. Therefore, T cell-mediated immune responses are thought to be crucial for combating this deadly pathogen. In chickens, gamma delta (γδ) T cells represent a major population with up to 50% of all peripheral T cells. However, their role in MDV pathogenesis and tumor formation remains poorly understood. To investigate the role of γδ T cells in MDV pathogenesis, we infected genetically modified chickens that lack γδ T cells (TCR Cγ^-/-^) with very virulent MDV. Strikingly, disease and tumor incidence were highly increased in the absence of γδ T cells, indicating that γδ T cells play an important role in the immune response against MDV. In the absence of γδ T cells, virus replication was increased by up to 89-fold in the thymus and spleen, both potential sites of T cell transformation. Taken together, our data provide the first evidence that γδ T cells play an important role in restricting MDV replication, pathogenesis and tumors caused by this deadly pathogen.

**Author Summary:** γδ T cells are the most abundant T cells in chickens, but their role in fighting pathogens remains poorly understood. Marek’s disease virus (MDV) is an important veterinary pathogen, causes one of the most frequent cancers in animals and is used as a model for virus-induced tumor formation. Our study revealed that γδ T cells play a crucial role in combating MDV, as disease and tumor incidence was drastically increased in the absence of these cells. γδ T cells restricted virus replication in the key lymphoid organs, thereby decreasing the likelihood of causing tumors and disease. This study provides novel insights into the role of γδ T cells in the pathogenesis of this highly oncogenic virus.

## Introduction

MDV is a highly oncogenic alphaherpesvirus that infects a wide range of chicken immune cells and causes deadly T cell lymphomas (1). Chickens are infected with the virus via inhalation of MDV-containing dust from a contaminated environment (2). In the respiratory tract, MDV is able to infect various immune cells including macrophages, dendritic cells (DCs) and B cells, which are thought to transport the virus to the primary lymphoid organs (3). The virus can be detected in the bursa of Fabricius, spleen and thymus within 24-48 hours post infection (3, 4). In these lymphoid organs, MDV infects B and T cells and subsequently establishes latency in CD4^+^ T cells (5). We could recently demonstrate that B cells are actually dispensable for MDV pathogenesis using the first cell-knockout chickens lacking B cells (6). MDV is also able to transform infected CD4^+^ T cells, which ultimately leads to deadly T cell lymphomas in various organs including liver, kidney and spleen. These tumors are mostly of clonal origin (7–9), indicating that only one or few T cells are transformed. Infected lymphocytes also transport the virus to the skin, where infectious virus is produced in the feather follicle epithelium (FFE) and shed into the environment (10). MDV infection can trigger both innate and adaptive immune responses. Various cell types are thought to be involved in the immune response against MDV including macrophages, natural killer cells, CD4^+^ and CD8^+^ T cells (11–13).

T cells are characterized by their T cell receptor (TCR), which can be divided into two main subgroups; alpha beta (αβ) and gamma delta (γδ) T cells (14). γδ T cells belong to the unconventional T cells and represent up to 50% of the peripheral T cells in chickens (5). The diversity of their TCR repertoires is greater than that observed in humans and mice (15). γδ T cells also represent a major subset of cytotoxic lymphocytes that can spontaneously lyse target cells without being restricted to MHC molecules (15). Until now, the role of γδ T cells in the immune response against many pathogens remains poorly understood.

Intriguingly, it has been recently shown that γδ T cells are significantly increased in MDV infected animals (5, 16). In addition, these cells up-regulate the expression of interferon-γ (IFN-γ) early in infection, suggesting they may play a role in either the immune response against MDV or its pathogenesis (16). Furthermore, it was recently shown that PBMCs activated with an anti-TCRγδ monoclonal antibody increase IFN-γ production and showed cytotoxic effect against MDV infected cells (17). An adoptive transfer of these PBMCs containing activated γδ T cells reduced virus replication in the lungs and MDV-induced tumorigenesis in chickens. This suggested that activated γδ T cells may play a role in initiating immune responses against MDV during the early stages of infection (17).

Despite recent advances, the role of γδ T cells in MDV pathogenesis remains poorly known, which is mostly due to the lack of γδ T cell-knockout chickens. Recently, we successfully generated a chicken line that lacks the γδ T cells (TCR Cγ^-/-^) (18). We used these knockout chickens to study the role of γδ T cells in the MDV life cycle. Our data revealed that the absence of γδ T cells increases virus replication in the thymus and spleen during early infection. In addition, we observed a drastic increase in both disease and tumor incidence in infected animals. Our experiments thereby shed light on the role of these abundant T cell population in the MDV pathogenesis.

## Results

### Absence of γδ T cells increases disease and tumor incidence

Until now, the role of γδ T cells in the immune response against MDV and its pathogenesis remains poorly understood. Therefore, we infected genetically modified chickens that lack γδ T cells with very virulent MDV. This chicken line was recently generated and characterized (18). Over the course of the infection, the disease incidence was significantly increased in TCR Cγ^-/-^ compared to the WT animals (Fig. 1A). Until the end of the experiment, 70% of the infected TCR Cγ^-/-^ animals showed MDV-specific clinical symptoms compared to 37.5% of their WT hatch mates. Similarly, tumor incidence was increased by more than 2-fold in the absence of γδ T cells (45%) compared to WT (20%) (Fig. 1B), suggesting that γδ T cells play a protective role in MDV pathogeneses. To decipher if the absence of γδ T cells affects tumor dissemination, the number of tumor-containing organs per tumor-bearing animal was determined. Surprisingly, the average number of tumors in the infected TCR Cγ^-/-^ animals was comparable to WT (Fig. 1C), suggesting that γδ T cells do not restrict tumor dissemination once tumors arose. Taken together, our data revealed that disease and tumor incidence is increased in the absence of γδ T cells, indicating that these cells play an important role in MDV pathogenesis and/or the immune response against the virus.

**Fig 1.**
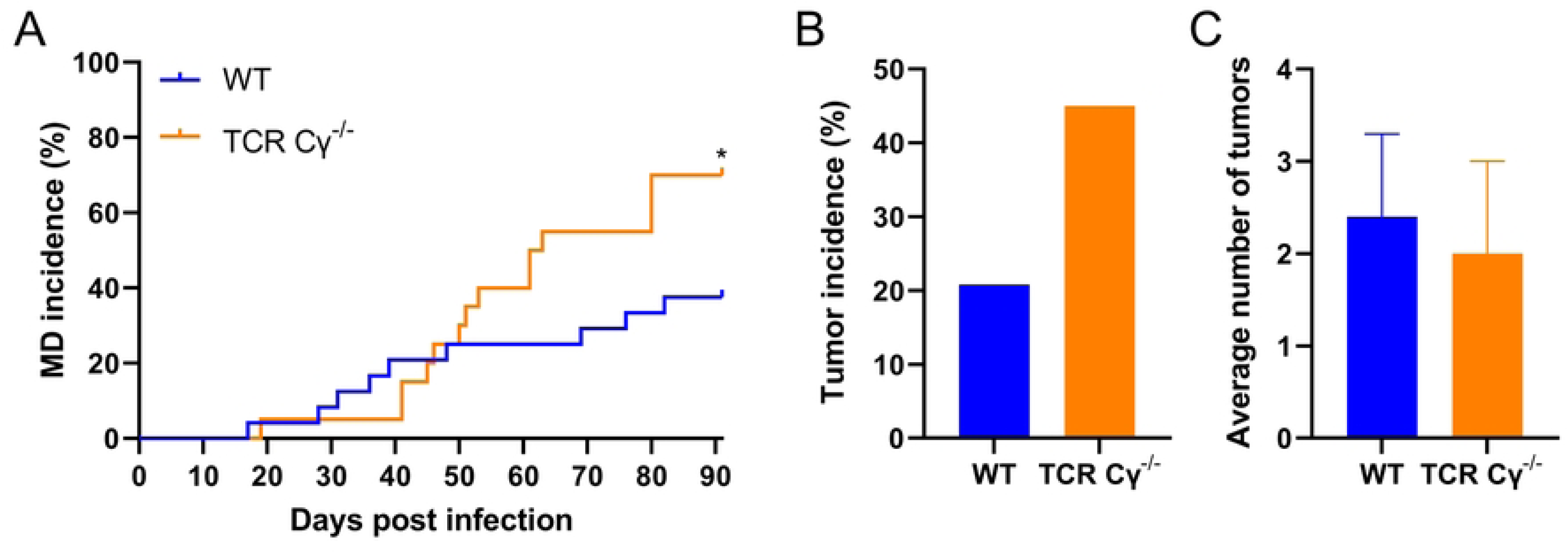
Absence of γδ T cells increases disease and tumor incidence. (A) Disease incidence in MDV infected WT (n= 24) and TCR Cγ^-/-^ chickens (n= 20). Percentage of chickens with clear clinical symptoms of Marek’s disease is shown over the course of the experiment (*p*<*0.05, Fisher’s exact test). (B) Tumor incidence is shown as a percentage of the chickens that developed tumors until the end of the experiment (*p*>*0.05, Fisher’s exact test). (C) The average number of gross tumor-containing organs per tumor-bearing animal is shown with standard deviation (error bars) (*p*>*0.05, Fisher’s exact test). Asterisks indicate statistical significance.

### γδ T cells are dispensable for MDV shedding and transmission to naïve birds

As γδ T cells have a high frequency in the skin (19), we investigate the role of γδ T cells in controlling virus replication in the skin, shedding and transmission. To achieve this, we quantified the MDV genome copies in feather shafts, dust and the infection of contact animals. Intriguingly, MDV genome copies in the FFE of TCR Cγ^-/-^ animals were comparable to WT animals (Fig. 2A), suggesting that γδ T cells are not involved in controlling MDV replication in the skin.

**Fig 2.**
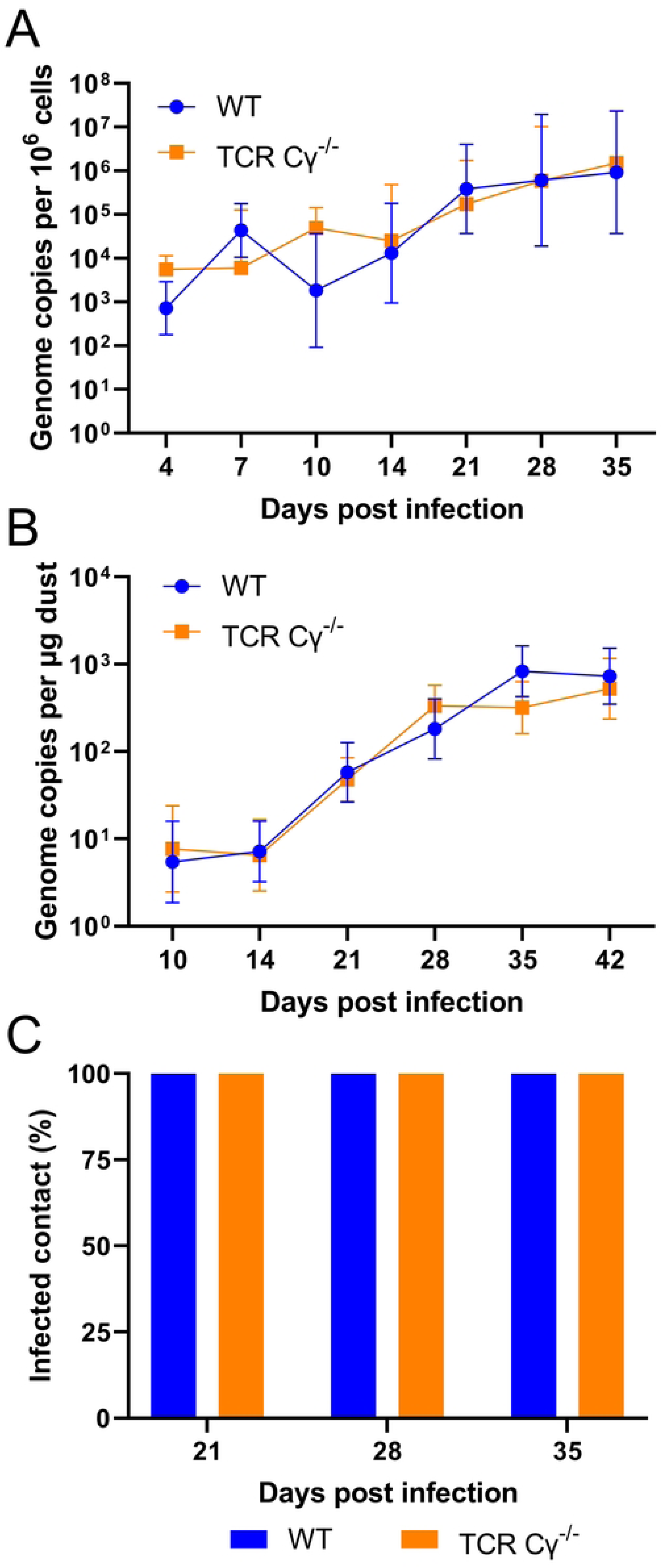
γδ T cells are dispensable for MDV shedding and transmission. (A) qPCR analysis of MDV genome copies in the FFE of WT (n= 8) and TCR Cγ^-/-^ chickens (n= 8). Mean genome copies are shown per million cells with standard deviation (error bars) (p*>*0.05, Mann-Whitney U tests). (B) Average MDV genome copies per 1µg of dust collected from the dust filter from each group at the indicated time points (20) (p*>*0.05, Mann-Whitney U tests). (C) Percentage of MDV positive contact chickens (n=8) detected by qPCR at the indicated time points.

Next, we evaluated the virus load in the dust. Consistently, MDV genome copies in the dust were comparable between both groups (Fig. 2B), indicating that γδ T cells do not influence virus shedding. In addition, we assessed if the absence of γδ T cells affects virus transmission. As MDV is efficiently shed into the environment after 14 dpi, we quantified MDV genome copies in the contact animals 21, 28 and 35 dpi (Fig. 2C). MDV was very efficiently transmitted to the naïve animals as all tested animals were already positive at 21 dpi. A comparable virus load was detected between the groups (data not shown). Taken together, these data reveal that γδ T cells present in the skin do not restrict MDV replication in the FFE, shedding and transmission.

### Impact of the absence of γδ T cells on MDV replication and immune cell populations in the blood

To determine why the disease and tumor incidence was increased in the absence of γδ T cells, we quantified virus replication in the blood at various time points. Surprisingly virus replication was comparable between the two groups (Fig. 3A), indicating that γδ T cells do not affect MDV replication in the blood. To determine if the absence of γδ T cells affects other lymphocyte populations, we quantified different populations including B cell, CD4^+^ and CD8^+^ T cells in the blood of the infected and uninfected groups on 7, 10 and 14 dpi. B cell numbers were not significantly different between the groups (Fig. 3B). The recently described decrease in the number B cell at 10dpi was observed in both infected WT and TCR Cγ^-/-^ birds (21). In addition, more B cells were detected in infected and uninfected TCR Cγ^-/-^ chickens at 14 dpi. Similarly, CD8^+^ αβ T cell numbers were also not statistically significant different (Fig. 3C), but again an increase was observed in infected and uninfected TCR Cγ^-/-^ chickens at 14 dpi. No significant differences were found for numbers of CD4^+^ αβ T cells (Fig. 3D), however, at 14 dpi we found an increase only in infected TCR Cγ^-/-^ animals. As MDV commonly transforms CD4^+^ T cells, this increase likely represents expanding tumor cells consistent with the increased tumor incidence in these chickens. Overall, this data highlights that γδ T cells do not influence the viral load in the blood and only had a minor effect on other immune cell populations in the blood.

**Fig 3.**
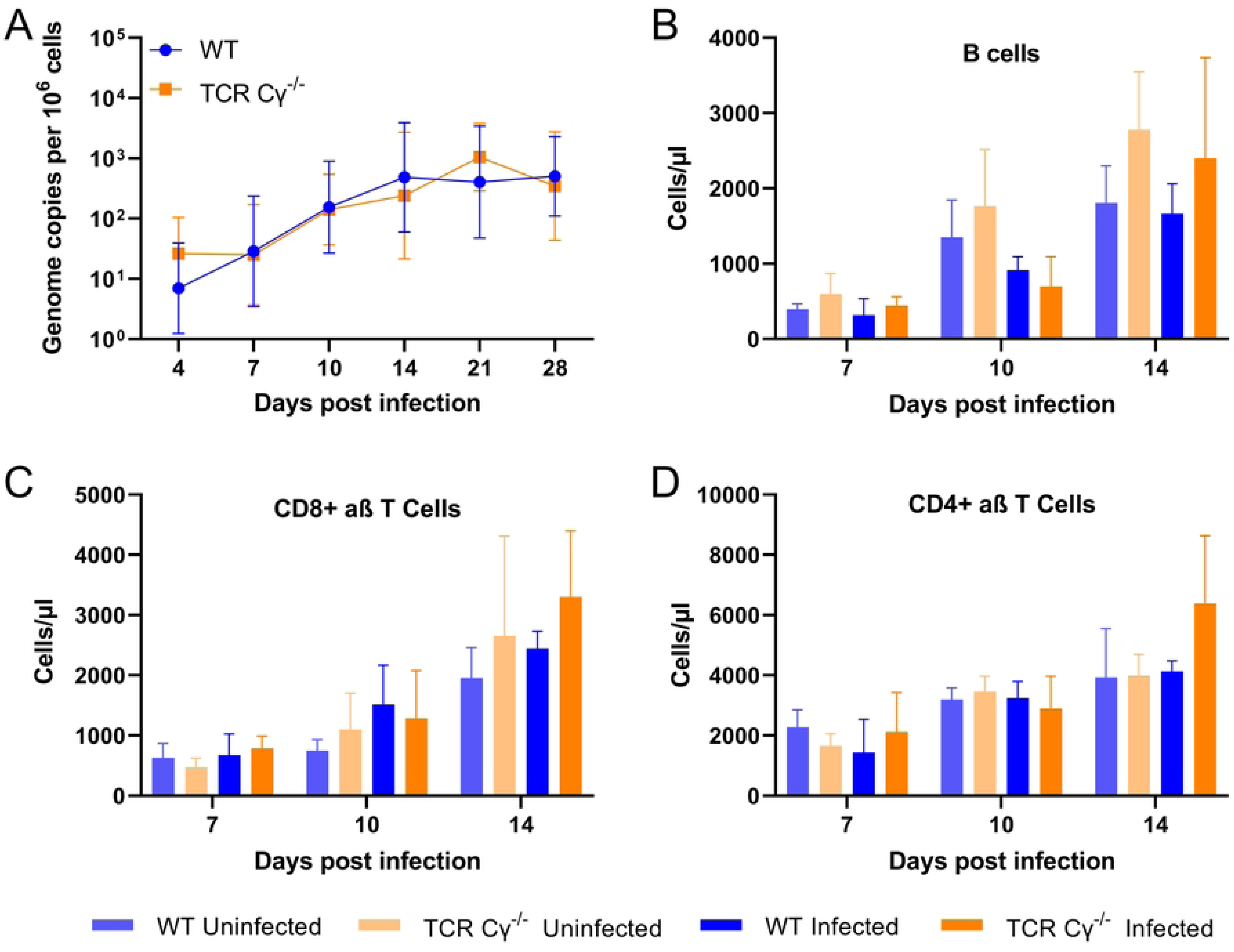
Impact of the absence of γδ T cells on MDV replication and immune cell populations in the blood. (A) qPCR analysis of virus replication in the blood of the infected WT (n= 8) and TCR Cγ^-/-^ (n= 8) chickens (p*>*0.05, Mann-Whitney U tests). B cell (B), CD8^+^ (C) and CD4^+^ T cell (D) count in the blood of uninfected and infected chickens WT (n= 3) and TCR Cγ^-/-^ (n= 3) using FACS (22) (p*>*0.05, two-way ANOVA (Tukey’s multiple comparisons tests)).

### Absence of γδ T cells increases MDV replication in specific lymph organs

To determine the role of γδ T cells in MDV replication in the primary lymphoid organs, we infected WT and TCR Cγ^-/-^ animals and quantified MDV genome copies in the bursa, spleen and thymus by qPCR. In the all three organs, comparable MDV genome copies were detected at 7 dpi (Fig. 4A-C), indicating that γδ T cells are dispensable for the delivery of the virus to the lymphoid organs. In the bursa which contains mostly B cells, a comparable viral load was detected during the phase of lytic MDV replication. Intriguingly, viral load in the spleen and thymus was markedly increased in the absence of γδ T cells at 10 and 14 dpi (Fig. 4B-C). These higher infection levels could increase the likelihood of T cell transformation and contribute to the elevated tumor incidence observed in the absence of γδ T cells.

**Fig 4.**
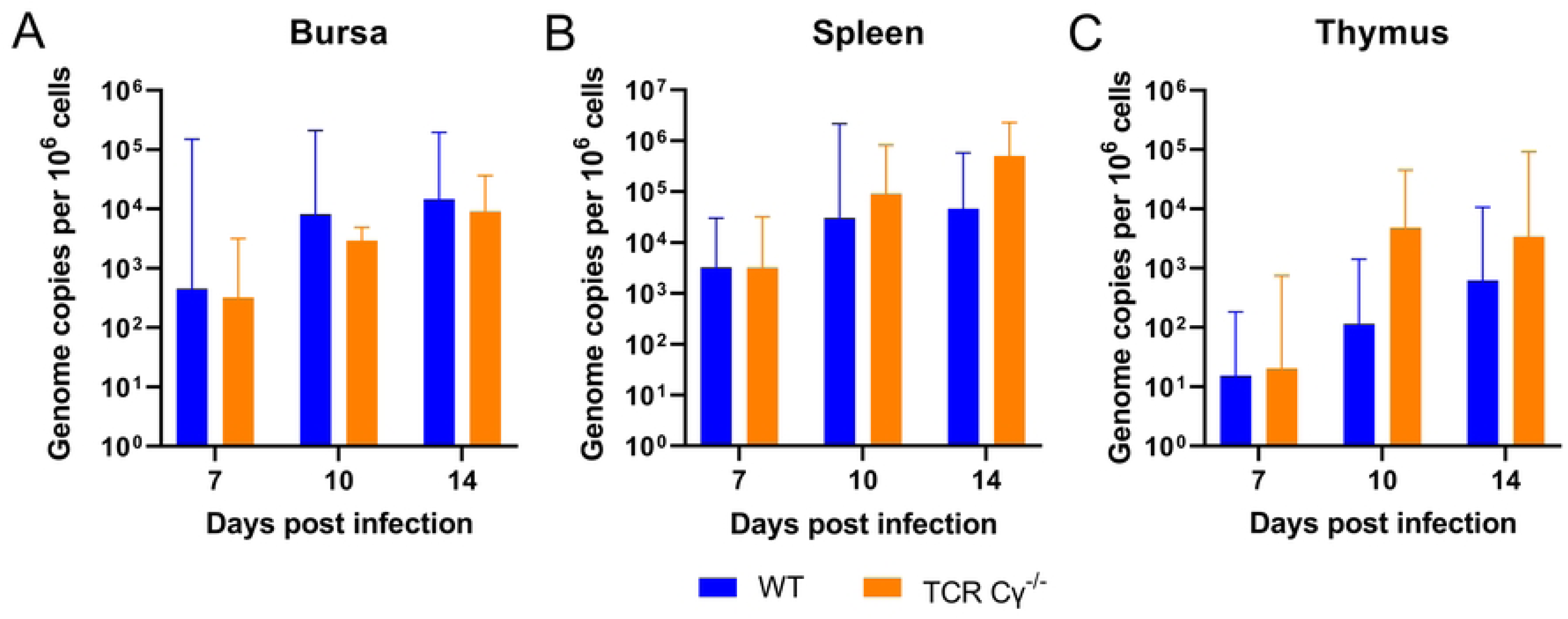
Absence of γδ T cells increases MDV replication in specific lymph organs. MDV genome copies in the bursa (A), spleen (B) and thymus of MDV-infected WT (n= 9) and TCR Cγ^-/-^ (n= 8) chickens at 7, 10, and 14 dpi. Mean genome copies are shown per million cells with standard deviation (error bars).

## Discussion

γδ T cells play a crucial role in the immune response against viral infections in mammals (23, 24). They possess the ability to recognize and kill pathogens and tumor cells in an MHC-independent manner (25, 26). In humans, γδ T cells have a frequency of about 5% of circulating T cells. In contrast, γδ T cells represent up to 50% of T cells in the blood of chickens (15, 27). A recent study revealed that γδ T cells can spontaneously trigger cytotoxicity to kill virus infected cells (15). Due to the highly cell associated nature of MDV, cellular immune responses in general are thought to be crucial to combat the virus. A recent study suggested that γδ T cells are likely involved in the immune response against MDV (17), a link that we followed up in our manuscript.

To investigate the role of γδ T cells in MDV pathogenesis and tumor formation, we infected chickens that lack γδ T cells with very virulent MDV (RB-1B strain) as suggested by Matsuyama-Kato et al (17). This recently generated and characterized chicken line allowed us to address the role of γδ T cells in MDV pathogenesis. In our experiment, we observed that in the absence of γδ T cells, the disease incidence was significantly increased during the course of the experiment. As tumors play a crucial role in the development of Marek’s disease, we determined if and how many infected knockout and WT animals developed tumors. Tumor incidence increased by more than two-fold in the absence of γδ T cells (45%) compared to the WT group (20%). This is relatively low for a virulent MDV strain and is due to the high genetic resistance of the chicken line (LSL, white leghorn) against MDV (18).

A recent study reported a delay in MDV tumor formation when PBMCs activated with an anti-TCRγδ monoclonal antibody were transferred into chickens. The study suggested that this delay is due to the upregulation of cytotoxic activity which could restrict MDV reactivation (17). In humans, γδ T cells were reported to have anti-tumor function against several types of lymphoma (28–30), and serve as a promising cancer immunotherapy.

Interestingly, the average number of visceral organs with gross tumors was comparable between TCR Cγ^-/-^ and WT animals. This suggests that γδ T cells do not restrict metastasis but only tumor development at an earlier stage.

It is known that infected T cells can transport the virus to the skin, where MDV efficiently replicates in the FFE and is shed into the environment (7, 31). Since γδ T cells have a high frequency in the skin (19), we investigated if the absence of these cells affected virus shedding. We quantified virus genome copies in the FFE, dust, and in naïve contact chickens. Surprisingly, comparable virus genome copies were detected in the feathers and dust of TCR Cγ^-/-^ and WT chickens by qPCR. This highlighted that γδ T cells do not influence MDV replication and shedding from the FFE. In addition, MDV efficiently spread independent of the presence or absence of γδ T cells as all contact chickens were infected until day 21 of the experiment. These contacts were all wild type chickens to ensure a comparable susceptibility to infection. The observation that virus genome copies were comparable between the groups indicates that comparable virus levels infected them in the same time frame. This is in agreement with a recent study which showed that MDV replication in the skin is not influenced by the infusion of PBMCs activated with an anti-TCRγδ monoclonal antibody (17).

To assess why TCR Cγ^-/-^ animals showed a higher disease and tumor incidence, we initially quantified virus replication in the blood of the infected animals over time. Intriguingly, the viral copies in the TCR Cγ^-/-^ animals were comparable to WT, suggesting that γδ T cells are dispensable for virus replication in blood. In addition, we assessed the effect of the absence of γδ T cell on other immune cell populations in infected and uninfected animals at 7, 10 and 14 dpi. B cell populations were not significantly different between the groups (Fig. 3B). Only slightly more B cells were detected in infected and uninfected TCR Cγ^-/-^ chickens at 14 dpi. CD8^+^ αβ T cell numbers were also not statistically significant different (Fig. 3C), while an increase was observed in infected and uninfected TCR Cγ^-/-^ chickens at 14 dpi. This is consistent with a previous study of von Heyl et al. that extensively characterized lymphocyte subsets in the blood of uninfected TCR Cγ^-/-^ animals and did not observe any significant changes compared to their WT hatch mates (18). Similarly, CD4^+^ T cells were also not significantly different (Fig. 3D), while only an increase in infected TCR Cγ^-/-^ was observed at 14 dpi. Since CD4^+^ T cells are the primary target for MDV transformation (3, 32), this increase may be due to the expansion of tumor cells.

Next, we assessed the role of γδ T cells in MDV lytic replication in the bursa, thymus and spleen. This is particularly important, as MDV mostly replicates in these lymphoid organs and transformation is thought to occur in them. In general, the virus was efficiently transported to the lymphoid organs as comparable levels were observed at 7 dpi, a commonly used time point for lytic replication. This indicated that γδ T cells do not play a role in the delivery of the virus to the primary lymphoid organs. Absence of γδ T cells did not affect virus replication in the bursa, likely due to the fact that the bursa is mostly composed of B cells and only few γδ T cells are present in the bursa that could affect MDV replication. Intriguingly, MDV replication was increased in the spleen and thymus in the absence of γδ T cells. The increase in virus load could have various reasons: it could be due to i) the lack of the γδ T cells that could combat the virus with their cytotoxic activity (5, 15), ii) an increase of CD4^+^ αβ T cells in infected TCR Cγ^-/-^ animals as they are the main MDV target cells of latency and tumorigenesis or iii) changes in organ structure and/or the distribution of immune cells in these organs in the absence of γδ T cells. Von Heyl et al. extensively characterized these organs but did neither observe changes in the immune cell populations (incl. CD4^+^ αβ T cells), nor changes in organ structure or the immune cell distribution in TCR Cγ^-/-^ chickens (18). This suggests that rather the lack of γδ T cells (i) is the reason for the observed phenotype than the immune cell composition (ii) or the organ structure (iii).

The increased virus load in the spleen and thymus but not in the blood, skin, or bursa, indicated that γδ T cells play a tissue-specific role in the immune response against MDV. This is consistent with a previous study that showed that γδ T cells have cytotoxic activity in the spleen but not in the blood (15).

In conclusion, our study provides crucial evidence that γδ T cells play an important role in MDV pathogenesis. Our data revealed a higher disease and tumor incidence in the absence of γδ T cells in MDV-infected chickens. Much higher viral loads were detected in the spleen and thymus in the absence of γδ T cells, indicating that γδ T cells restrict virus replication and/or tumor development. Overall, our data provide important insights into the role of this highly abundant cell population in the pathogenesis of this deadly pathogen.

## Materials and Methods

### Ethics Statement

All animal experiments were conducted according to the relevant international and national guidelines for the humane use of animals. The permission to conduct these experiments was granted by the Landesamt für Gesundheit und Soziales (LAGeSo) in Berlin, Germany (approval number G0294-17 and T0245/14)

### Genotyping

Whole peripheral blood was collected from newly hatched chicks and total DNA was extracted using the NucleoSpin 96 Blood core kit (Macherey-Nagel, Düren, Germany) according to the manufacturer’s instructions. Genotyping has been performed by PCR using TCR-specific primers as published previously (18). Chicks were categorized into two groups: WT (TCR Cγ^+/+^) or KO (TCR Cγ^-/-^). Primers used for genotyping are in Table 1.

**Table 1.**
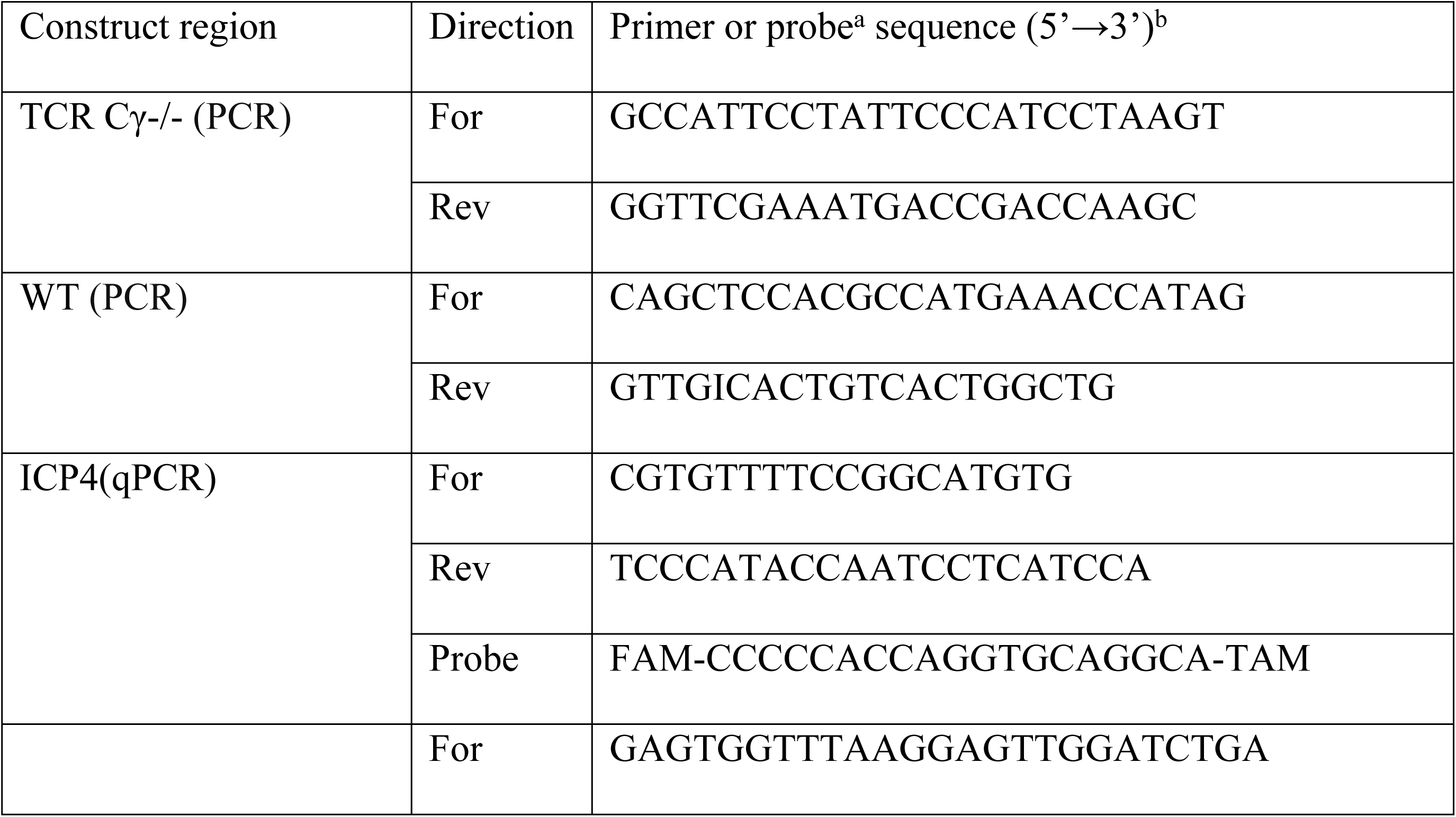
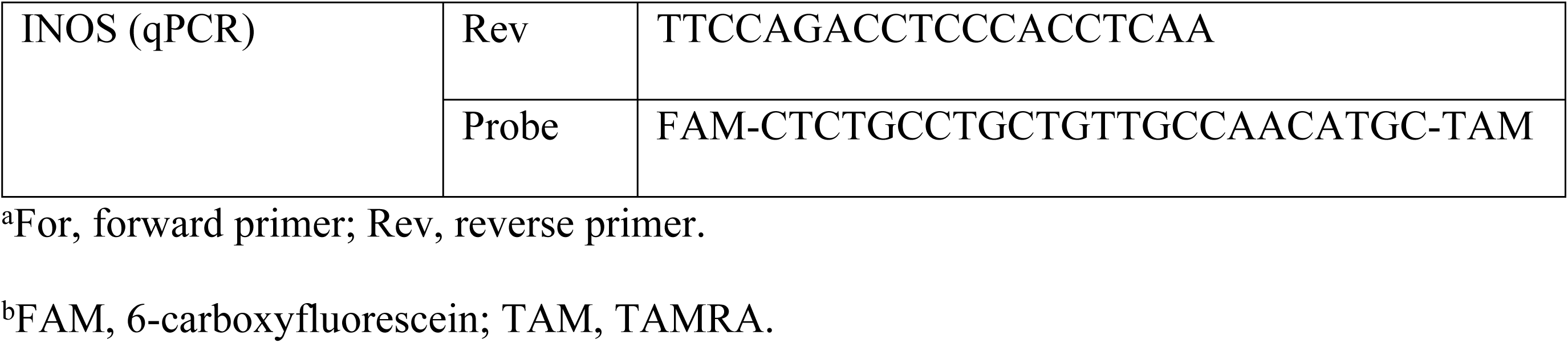
PCR and qPCR primers and probes used in this study.

### Cells and viruses

Chicken embryo cells (CECs) were prepared from 11-day old Valo specific-pathogen free (SPF) embryos (ValoBioMedia) as described previously (33). CECs were propagated in MEM (Pan-Biotech, Aidenbach, Germany) supplemented with 1-10% FCS, 1% penicillin/streptomycin at 37°C under a 5% CO2 atmosphere. The very virulent RB1B wild-type strain was propagated on CECs, stocks were frozen in liquid nitrogen and titrated prior to their use.

### Animal experiments

#### Animal experiment 1

To investigate the role of γδ T cells in MDV-induced pathogenesis, one-day-old chicks were genotyped. Wild type (WT; n= 24) and γδ T cell knock out (TCR Cγ^-/-^; n= 20) animals from the same parents were infected subcutaneously with 2000 PFU of very virulent RB-1B strain. To assess the natural transmission of the virus, one-day-old VALO SPF (VALO BioMedia) chickens (n=11 per group) were housed with the infected ones. The two groups were housed separately and supplied with food and water *ad libitum*.

To assess virus replication in the infected animals, peripheral blood was collected at 4, 7, 10, 14, 21 and 28 days post-infection (dpi). To quantify the virus genome copies in the skin of the infected animals, feather samples were collected at 4, 7, 10, 14, 21, 28 and 35 dpi. To quantify the shedding of MDV into the environment, dust was collected in the rooms at 10, 14, 21, 28, 35 and 42 dpi. To assess the infection of the contact animals, peripheral blood was collected at 21, 28 and 35 dpi. Chickens were monitored on a daily basis throughout the experiment for the development of MDV-specific symptoms including ataxia, paralysis, torticollis, and somnolence. Once chickens exhibited severe symptoms or at the end of the experiment (91 days), they were humanely euthanized and examined for gross tumors and the spleens were collected to assess the virus load.

#### Animal experiment 2

To determine if the absence of γδ T cells affects virus replication in the lymphoid organs, one-day-old chicks were genotyped, divided into two groups WT (n=9) and TCR Cγ^-/-^ (n=8) and infected as described above. In parallel, uninfected control chickens (WT; n = 9, TCR Cγ^-/-^; n = 6) were raised in a separate room.

Blood samples were collected from infected and control animals at 7, 10, and 14 dpi. To assess the delivery to and replication in the lymphoid organs, MDV genome copies were quantified in the spleen, thymus and bursa at these time points.

### DNA extraction and genomic quantification of the virus

Whole blood DNA was extracted using the NucleoSpin 96 Blood core kit (Macherey-Nagel, Düren, Germany) according to the manufacturer’s protocol. DNA was also extracted from feathers and dust using a proteinase K lysis protocol described previously (20). DNA from organs was extracted using the innuPREP DNA mini kit (Analytik-Jena, Berlin, Germany) following the manufacturer’s instructions. To quantify the virus load by qPCR, specific primers and probes (Table 1) for MDV ICP4 were used. The virus genome copies were normalized against the chicken induced nitric oxide synthase (iNOS) gene (10, 34, 35).

### Flow cytometry

To assess the effects of infection and γδ T cell knockout on other immune cell populations (incl. thrombocytes, monocyte, T and B cells), absolute counts of these cells in the blood were determined by flow cytometry as described previously (22). Briefly, the peripheral blood was collected in precoated anticoagulant tubes and stabilized with the TransFix ® reagent (Cytomark, Buckingham, UK) according to the manufacturer’s instruction. Whole blood was diluted with flow buffer, incubated with an antibody mix of anti-TCRαβ/Vβ1-FITC (clone TCR2) anti-TCRαβ/Vβ2-FITC (clone TCR3), anti-TCRγδ-PE (clone TCR1), anti-Bu1-Pacific Blue (clone AV20), all Southern Biotech, Birmingham, USA, anti-CD8-PerCP-Cy5.5 (clone CT8, Southern Biotech, LYNX Rapid PerCP Antibody Conjugation Kit, Bio-rad, Feldkirchen, Germany), anti-CD45-APC (clone UM16-6, LYNX Rapid APC Antibody Conjugation Kit, both Bio-rad) and thrombocyte marker K1-PE (LYNX Rapid RPE Antibody Conjugation Kit, Bio-rad) (36). Flow cytometric measurements were performed with a FACSCanto II (Becton Dickinson, Heidelberg, Germany) and data were analyzed using the FACSDiva (Becton Dickinson, Heidelberg, Germany) and FlowJo (FlowJo LLC, Oregon, USA) software (S1 Fig).

### Statistical analysis

Statistical analyses were performed using Graph-Pad Prism v9 (San Diego, CA, USA). The MD incidence graph was analyzed using the log-rank test (Mantel-Cox) test. Fisher’s exact test was used to assess the MD incidence at the final necropsy (91dpi). The tumor incidence and the average number of tumors per animal were analyzed using Fisher’s exact test. MDV genome copies in the feather or dust were analyzed using the Mann-Whitney U test. MDV genome copies in the blood of experimentally infected and contact animals were analyzed using the Mann-Whitney U test and paired t-test respectively. The immune cell counts were analyzed using the two-way ANOVA (Tukey’s multiple comparisons tests).

## Acknowledgments

We are grateful to Ann Reum for her technical assistance. Thanks to Lisa Kossak for her help in an animal experiment. Thanks to Dr. Roswitha Merle in the Institute of Veterinary Epidemiology and Biostatistics, Freie Universität Berlin, for the statistical assistant.

## Supporting information

**S1 Fig. Flow cytometry gating.** EDTA-blood was stained with mix of anti-TCRαβ/Vβ1-FITC), anti-TCRαβ/Vβ2-FITC (clone TCR3), anti-TCRγδ-PE (clone TCR1), anti-Bu1-Pacific Blue (clone AV20), anti-CD8-PerCP-Cy5.5 (clone CT8, anti-CD45-APC (clone UM16-6,) and thrombocyte marker in a no-lyse no-wash one-tube procedure and subsequently analyzed by flow cytometry. We first separated thrombocytes and leukocytes (A) followed by a single cell gate (B and D). Leukocytes were subdivided in B cells (E), yd-T cells (G) and aß-T cells (J). T cell subpopulations were further separated in CD8pos and CD8neg cells (I and L). CD8neg aß-T cells were addressed as CD4pos T cells. To exclude potentially contaminating erythrocytes, for all cell populations an additional FSC/SSC gating was performed (C, F, H and K).

